# The Cultural Evolution of Hard-to-fake Rituals

**DOI:** 10.1101/060640

## Abstract

It has been proposed that costly rituals act as honest signals of commitment to group beliefs when such rituals appear dysphoric and unappealing (costly) to non-believers, but appealing to true believers (Irons, 2001). If only true believers are willing to endure ritual behaviors and true belief also entails belief in altruistic cooperation, associating with other ritual practitioners can help solve cooperation dilemmas in groups by sorting out potential free-riders. While this hypothesis is obviously true if such ‘faking’ of ritual is strictly impossible, strict impossibility seems implausible. ‘Faking’ is defined by Irons in this context to be to be performing the ritual without commitment to group beliefs. In this paper, I posit various ways that such faking might be difficult, instead of impossible, or different ways in which such ritual faking might be ‘costly’ and then formally model the social learning and cultural evolution dynamics to see where it may still hold theoretically that such rituals help maintain altruism in group and under what conditions. Analytic solution for evolutionary equilibrium is derived for each model, verifying that under a wide range of conditions for some, but not all interpretations, such hard-to fake rituals can help groups solve cooperative dilemmas, including in some circumstances that might not be intuitively obvious, such as where such free-riding is not visible and free-riders successfully represent themselves as true believers to observers.

It is also the case that while there has been some progress in cleaning up the definitional confusions in the animal signaling literature around costly signaling, the literature on human rituals as costly signals has introduced novel uses of the term ‘cost’. Theories referring to completely different mechanisms or even definitions of ‘cost’ are sometimes conflated. To contextualize the analysis of costly-to-fake rituals, this paper provides a review of the ideas proposed in the literature on costly human rituals and differentiates them from costly signaling as used in the animal behavior literature.

## Introduction

Religions and other social groups often have ritual requirements that seem aversive to outsiders but create deep senses of meaning and belonging to group for participants. On the one hand there are the time consuming requirements to daily prayer that characterize many religions, as the 5 daily prayers required of Muslims in the Quran. On the other are more sensorially extreme practices like fire walking (Xygalatas et al., 2013) or the combinations of ritual piercing, fasting, and durational dancing and chanting characteristic of the Lakota Sun Dance (Peltier, 1999; Portman & Garrett, 2006). Such activities are puzzling to outsiders, and this obscurity of function is itself part of the definition of ‘ritual practices’ used by many students of cross cultural comparison (Durkheim, 1912). From an evolutionary perspective, the maintenance of costly activities without clear productive function is a puzzle. Put simply, wouldn’t groups that spent their energy more efficiently produce more wealth and come to be more successful, or wouldn’t those behaviors for which we have an aversion to learning or practicing disappear out of populations because of this aversion? Irons proposes that many religious or ritual activities are “hard-to-fake”, being disagreeable or insufferable to non-believers, but that when we adopt the belief system of the religion or group, the belief system shifts our perspective on and experience of the ritual, making it something desirable to do (Irons, 2001). They are hard-to-fake or costly-to-fake in that those who do not have true belief have incentives to not perform the ritual, some extra difficulties in doing so, or perhaps a lack of positive associations with it. The belief system changes our values and experience. To the non-religious, it may seem repugnant to pierce oneself, to walk across a bed of hot coals or to spend many hours regularly in ceremony or days in fasting, but to the devout, such practices can be meaningful, making up for or even reversing any potential negative experiences from them (Cruwys et al., 2013; Peltier, 1999). Serving as embodied markers of meaning and identity, such rituals may truthfully mark those with very strong feelings of altruistic commitment to group, willingness to sacrifice for the group, as when a small group of contemporary Sun dancers held off a group of several hundred Canadian police in an armed conflict over their sacred group practices and identities at Gustafson Lake, BC in 1995(Shrubsole, 2011). According to Irons’ theory, these rituals successfully serve as markers of altruistic intent because few but true believers would willingly perform the otherwise very costly ritual. Where prosociality is part of such a belief system, this difference in how ritual is experienced facilitates positive assortment of altruists. As ritual performance is an honest marker of commitment to the belief structure, engaging in cooperative relationships preferentially with co-ritualists will help avoid the problem of free-riders: those who take advantage of altruistic relationships without themselves returning the altruistic favor.

This hypothesis relies critically on an assumption that ritual practice is linked to true beliefs in altruism. It assumes that the belief system is difficult to parse apart or that it is difficult or aversive somehow to practice the ritual without having the group beliefs. The individual can not *easily* adopt just that part of the belief system that changes the perceived value of the ritual trait without adopting that other part of the belief system that calls on individuals to be altruistic toward co-religionists. Without such belief, the ritual practice appears or is experienced as boring, painful, or otherwise aversive without compensating spiritual benefits. This parallels asserting an intrinsic trait linkage in a genetic signaling model (Hamilton, 1964). While the hypothesis of accurate signaling of altruistic intent is by definition true if such separation is impossible, strict impossibility itself seems implausible. This is the central problem I address in this paper, whether such hard-to-fake rituals can help groups culturally evolve through solving free-rider problems when ritual is hard, not impossible, to fake. I posit various interpretations of how such parsing or faking of group beliefs might be *difficult*, rather than impossible. I then formally model social learning and cultural evolution dynamics to see where the hypothesis that hard-to-fake rituals serve as reliable markers of altruistic intent may theoretically still hold and under what conditions. I find that it does hold, but not always. In the simplest cases, where non-believers’ aversion to perform is greater than the cost of cooperative altruism, such ritual facilitated altruism is sustained in groups. A particularly interesting sub-example, from a theoretical perspective, is how fakers might falsely advertise true beliefs. They then act as free-riders, but also help the group beliefs spread in social learning by advertising true belief. I find that despite being a drain on groups, such selfish free-riders may actually benefit groups by acting as (false)advertisers of true belief. In this case, true belief may still move to fixation even though the cost of altruism is less than the experienced aversion to faking rituals. They will at first bolster groups that would otherwise dwindle through social learning, but then be eventually eliminated from the population themselves as groups became homogenous groups of ritual performers, with no overt models of non-belief. Further, when there is a possibility of costly investment in exposing free-riders, ritual practice will move to fixation with a balance of freeriders and true believers.

A faked ritual may be costly by being experientially dysphoric, cognitively difficult to innovate, hard to socially learn because it is veiled, or perhaps by having a genetic fitness cost. These different kinds of costs lead to different dynamics. Analytic solution for evolutionary equilibrium is derived for each model, verifying that under a wide range of conditions for some, but not all models, such hard-to fake rituals can help groups solve cooperative dilemmas. The point of this paper is not to prove that the psychological premise of the theory is empirically accurate in specific cases, but to more precisely formulate it, and then through evolutionary models, see if the premise leads to the conclusions of the theory. It would be an empirical question whether these psychological dispositions are good descriptors of human behavior in a given context. Certainly there is evidence that such is at least plausible (Irons, 2001; Sosis & Bressler, 2003).

## Disambiguation of the term *cost* in relationship to signaling theory in general and rituals as signals in particular

It has been noted that the notion of cost in relation to theories of signals or displays has been in a messy state, with very different arguments around the function of cost or even definition of what a ‘cost’ might be, combined with a fair bit of conflation of these arguments (Cronk, 2005). While some progress has been made in clarifying the differences in the various uses and arguments around ‘costs’ in relation to ‘signaling’ in the animal behavior literature (Maynard-Smith & Harper, 2003), the literature on humans and socially learned rituals presents some new uses of the term ‘cost’ and some very different arguments about such in relationship to rituals as signals. Some further disambiguation is in order before these theories can be evaluated for cogency.

Maynard Smith and Harper differentiate two different kinds of costs that are often referred to in theories of signals in animal behavior: *efficacy* costs and *strategic* costs. Costs here are understood as reductions in genetic fitness. It is the latter, strategic cost, that is the usual subject of ‘costly signaling theory’. An efficacy cost is a cost that is required mechanically in order to make the signal in a way that is perceived and reacted to by the receiver. This concept parallels Cronk’s notion of reception psychology (2005), where the idiosyncrasies of the existing evolved psychology or perceptual mechanics requires a cost for transmission. This is exemplified by a ‘manipulative signal’, where a signal takes advantage of a pre-existing perceptual structure or bias of the receiver to manipulate the behavior of the receiver, potentially to the detriment of the receiver (Krebs & Dawkins, 1984; Ryan, 1998). Another example would be an ‘indexical signal’, like stotting in antelope (FitzGibbon & Fanshawe, 1988), where a signal (like height of jumping) is only possible if one has the quality (strength) being signaled. Similarly, where coalition strength is important information to share with adversaries in order to avoid mutually costly combat, massing is an indexical signal of group size. In these cases, it may be worth the signaler pay a cost to make the signal, given the benefit of the signal’s reception. If such a cost is mechanically necessary, it is called an ‘efficacy cost’. Efficacy costs are not necessarily associated with conveying information for the consideration of a receiver, but manipulate a behavioral tendency or sensory bias of the receiver. If an identical signal were possible without the cost, evolution would favor the costless signal. A strategic or handicapping cost, however, is one where the cost is only worth paying if one has the quality or a sufficient amount of the quality. This is the usual subject of ‘costly signaling’ models in the animal behavior literature, where the cost itself is necessary for the signal to function (Grafen, 1990; Zahavi, 1975). An individual of lesser quality will find that the cost of the signal is not worth the advantage to be gained from the signal and so will not pay the cost. In contrast to the case of an efficacy cost, if an identical signal were introduced that was costless, then the individuals lesser in quality would make the signal to get the benefit from the receivers, the receivers would stop paying attention to the signal since it no longer carried useful information, and signaling would then disappear from the population. Strategic handicaps/costs guarantee the information content of a signal to which the receiver responds strategically. Versions of the theory of strategic costly signaling have been successfully used to explain, in humans, a wide range of costly behavior, including costly and inefficient hunting behavior (Hawkes, 1991; Smith, Bliege Bird, & Bird, 2003), growing inedible but conspicuously large yams (Bliege Bird & Smith, 2005), and the conspicuous consumption and expensive fashions of the wealthy (Veblen, 1899).

Key for the purposes of this paper is that such strategic costs do not signal intent. They signal quality and intent is assumed to be obvious. Costly signaling in this sense can not be used to explain the intent to perform altruistic behavior. It may in some cases be a useful explanation of *conspicuous* altruism when it itself is used *as* a costly signal (Gintis, Smith, & Bowles, 2001), but it is not itself an explanation of a hard-to-fake signal of intent to be altruistic, especially where such altruism is not visible to the recipient. The strategic cost only guarantees a specific level of some quality in the signaler and does not constrain their response to the receiver’s reaction. It *might* signal a resource capacity to be altruistic, but not an intention to be altruistic. While it is empirically observed that costly rituals are associated with effective group bonding and in-group altruism (parochialism) in humans (Ruffle & Sosis, 2007; Sosis & Bressler, 2003), the cause of this association would not in any way be explained through the dynamics referred to in the literature on costly handicap signaling in animals (Grafen, 1990; Sosis, 2006; Zahavi, 1975), and the analysis of costly signaling in these contexts likewise can not simply be extrapolated as support for theories of costly signaling of intent.

In the literature on human rituals, a number of additional proposals for the function of cost in ritual are in use. While these ideas of costly signaling are sometimes conflated or assumed to be related phenomena (Sosis, 2006), they are mechanistically different, unrelated proposals. These hypotheses include 1) rituals as hard-to-fake signals, 2) sacrifice and stigma as costly boundaries for maintaining contribution to public/club goods, 3) bonding rituals as hijackers of pre-existing behavioral dispositions, 4) runaway cultural evolution processes, 5) rituals as ethnic markers and shared norms. Below, these distinct theories of cost and ritual are defined in more detail and differentiated. I use cultural evolution modeling to analyze the claims of the hard/costly-to-fake theory of ritual. The analysis does not bear directly on these other theories of cost in relation to signaling. They are reviewed and clearly disambiguated because the conflation of these theories has significant potential to generate confusion about what should be considered distinct and unrelated social mechanisms. In other words, the intention is not to relate them, but to show them as unrelated in order to clarify the discussion.

### Rituals as hard-to-fake signals (Irons, 2001)

As described in the introduction, Irons presents a verbal argument for human rituals as ‘hard-to-fake’ signals of altruism; rituals which are costly to non-believers but cheap to believers signal honest belief in a group’s belief structure, and when altruism is an essential part of the belief system, they thereby honestly signal altruistic intent. The word ‘cost’ is being used in a very different way than in the animal signaling literature, and what is signaled is not quality, but intent. Cost here refers to the psychological experience, rather than to effects on genetic reproduction. What is signaled is not some level of a quantitative capacity of the signaler, but intentions of the signaler. As such, this has more in common with a Green Beard model than with a costly signaling model.

This verbal model is a cultural evolution model, not a genetic one. To ‘fake the ritual’ would be to either choose to endure suffering that true-believers are not experiencing or to find some other hard-to-acquire technique for the transformation of the experience besides the belief structure of the group. There would be either an aversion to copy the former or a difficulty in acquiring the latter. Examples might include high arousal rituals involving very strong sensations, like firewalking, whipping oneself (Christian flagelentes) or ritual piercing (Sundance) or low arousal rituals with long time commitments, like praying 5 times per day, attending a long weekly sermon, or performing regular religious offerings (puja) (Atkinson & Whitehouse, 2011). The experience or time investment of these rituals seem unpleasant for most non-believers, but to believers are a potentially ecstatic experience based on mental/spiritual commitments. While it might not incur a significant reproductive cost (in comparison to the advantages of free riding on group membership), the sensation prevents most non-believers from being able to do it, whether it be the intensity of strong sensation or the boredom of long time investment. However, ‘true belief’ transforms the sensation into a kind of ecstasy, a transformation of the sensation which others might not find possible or easy to do without belief. Faking the ritual is possible but very difficult to learn. It is an odd assertion, but it is certainly true that beliefs can change reaction to or perception of sensation (Xygalatas, Konvalinka, Roepstorff, & Bulbulia, 2011), and it is at least a plausible hypothesis that it may in some circumstances be difficult due to pre-existing cognitive/emotional biases to selectively copy those aspects of a group’s beliefs that transform the sensation without also acquiring the group’s beliefs that cause self enforcing of altruism. For example, acquiring the whole package of beliefs may cause less intrapsychic costs and hence less damaging stress levels or a lower risk of making a mistake in social relations than picking and choosing. In fact, subtle aspects of expression may be out of conscious control, being based in the limbic system, and thus proper ritual performance may actually be impossible without a shift in belief (Eckman, Levensen, & Freisen, 1983). Of course, those who are part of the BDSM (“Bondage, Discipline, Sado-Mashocism”) community are also capable of experiencing flagellation and prostration as pleasurable outside of authentic religious contexts (Zussman & Pierce, 1998), so such the theory would require that selfish masochists would be rare. Using cultural evolution models, I demonstrate below how groups may use such costly or hard to fake rituals to support in-group altruism, despite some amount of free-riding fakers. I also show the constraints on this ability and the conditions under which such cooperative groups are overcome by free-riders.

It is important, as we try to model a specific costly ritual, that we not simply assume that a painful experience is necessarily disfavored by social learning. This may be true, but the potential negative experience may also cause the behavior to be more strongly noticed by an observer, which may paradoxically lead to a positive social learning bias (Nuttin, 1975).

Note, Henrich has addressed a related phenomena with the Credibility Enhancing Displays (CREDs) model (Henrich, 2009). In the verbal set up of the CRED model, true beliefs are similarly invisible. We have a tendency to look for costly displays as evidence of true beliefs, and when we see something that we *believe* is a CRED in an observed other, we are more likely to acquire the expressed belief of the other as a true belief ourselves. In the costly-to-fake ritual verbal argument, it is proposed that certain ritual types are likely to be associated with true belief. Such hard-to-fake rituals would be *valid* CREDs of true belief, although Irons does not suggest in his verbal model that we learn true belief based on them. Irons simply claims that they can be successfully used functionally to evaluate belief of others and filter out free-riders. Likewise, the CRED model does not look at whether or not the CRED psychology of social learning biases could itself evolve in the context of Public Goods and free-rider problems. It answers what happens if we believe something is credibility enhancing, but not what we find credibility enhancing and what should, evolutionarily speaking, be credibility enhancing. There needs to be more empirical work here on the circumstances in which CREDs and CRED psychology operates and the form it takes in relation to cost, as well as modeling work on when should it operate in the context of PG dilemmas and free-riders. While it is not the purpose of this paper, synthesizing these two questions would be a useful modeling project.

### Sacrifice and stigma as costly boundaries that maintain contribution to club goods (Iannaccone, 1992)

Iannaccone, using microeconomic theory, models individuals as rational optimizers facing a problem of club goods. Religious groups are considered as providers of *club* or *public* goods which are greater with greater per capita participation in the group. Such goods are non-excludable to group members, but excludable to non-group members, and there is more available as individuals within the group altruistically contribute to the good. Ritual participation is viewed here as stigmatizing, preventing participation in out-group economic activities, thus requiring participation in sect-structured activities which are unavoidably ‘altruistic’. The ‘cost’ of the theory is a different take on cost, being this stigma or barrier to productive interaction outside the group. The stigma means that people are either ‘all-in’ or ‘all-out’ and prevents partial participation, which could manifest as free-riding. This stigma is necessary, as such free-riders might otherwise engage in economic activity for their own benefit outside of the group, but then take advantage of the profits of group industry while only minimally participating in generating these goods. The stigma prevents or reduces such investment outside of the group. While this can account for very specific forms of public goods contributions, this model, does not account for unenforced altruism within the group. An assumption of the model is that full group participation absolutely requires contribution to the public goods in question, there not being structurally a possibility to free-ride. It does not account for uncoerced cooperation within the group. It may account for such things as participation in church industrial activities, like Shaker furniture making, but does not account for such things as doing one’s dishes when no one is watching or engaging in such manufacture as well as one can. Where contributing to the public goods of the group is not coerced, lannacone’s model will not support altruism.

In this model, the ritual participation is an indexical display of stigma and thus an honest signal of structural constraints on the individual’s behavior. Their quality is irrelevant to the model, and intent does not need to be known since action is constrained by the social structure. The hard-to-fake hypothesis rests on the proposal that it solves a problem of altruistic action within groups. Iannaccone’s model assumes that if people are constrained from acting outside of the group, they will intrinsically contribute altruistically to the club goods. It assumes away a problem that the hard-to-fake proposal addresses, and it is this problem which I investigate in this paper.

### Bonding Rituals as hijackers of pre-existing behavioral dispositions (Frost, 2016a, 2016b)

In this model, a reciprocal ritual practice is innovated which hijacks an existing behavioral predisposition which induces altruistic behavior toward the ritual co-participant. This is similar to Dawkins and Krebs argument of costly signals taking advantage of pre-existing sensory biases. However, here it is not just the other that is manipulated but the self, and the ritual, being reciprocal, can only be done with a partner also doing the ritual. This existing predisposition is assumed to have a genetic basis, and evolutionary models demonstrate that in a wide range of circumstances, but not all, this can lead to ritual performance and altruistic response at evolutionary equilibrium. Specifically, where ritual performance is introduced to a population as either a genetically based or socially learned behavior, coevolutionary dynamics may lead to any combination of ritual performance or not, altruistic response or not, depending on the relationships of ritual cost, benefit and cost of implied public goods and relative a priori benefit of the preexisting hijacked genetic allele. In the purely genetic model, the cost of the ritual is a genetic fitness penalty. In the gene-culture coevolution model it is both a genetic fitness penalty and a bias against social learning proportional to the fitness consequences, via success-biased social learning. For a significant range of parameter values, the model demonstrates long term cycling of alleles in the population.

There are a number of specific instantiations of this general dynamic which are supported empirically. The research on synchronous movement (McNeill, 1995; Reddish, Fischer, & Bulbulia, 2013) and unconscious mimicry (Chartrand & Baaren, 2009) strongly suggests that rituals involving such synchrony triggers prosocial instincts. The dual modes of religiosity hypothesis (Whitehouse, 2002), suggests two ritual forms: *imagistic* mode rituals are those that are high arousal, dysphoric, and infrequently performed, bonding small groups tightly, while *doctrinal* mode rituals are those that are low arousal, regularly recurring dysphoric rituals and cause lighter bonds in much larger groups. This is supported empirically, through systematic assessments of the associations of rituals forms and polity size in the ethnographic record (Atkinson & Whitehouse, 2011). Finally, calming practices like meditation have been shown to increase altruism in groups and decrease parochialism (Frost, 2013), and when such altruistic practices are associated with positive assortment amongst coreligionists, such would also create in-group altruism (Wilson, 2002).

This theory of ritual hijacking of pre-existing sensory biases has a lot of similarities to Irons’ costly-to-fake rituals. Both involve ritual activities that allow assortment amongst ritualists and involve a linking of ritual activity and parochial altruism that potentially solves free-rider problems, being a signal of intent and not quality. The underlying psychological mechanism linking altruism and costly ritual performance is different. I look at the evolutionary dynamics of such mechanisms in a separate paper (Frost, 2016b)

### Social learning of learning biases and runaway cultural evolution processes (Boyd & Richerson, 1985)

It has been shown that where a bias in social learning is itself socially learned, there can be a runaway effect where the behavior which the bias favors and the strength of the bias may evolve together in a runaway process, potentially leading to quite exaggerated traits. Such exaggerated traits could be costly in terms of other instinctual learning biases based on the genetic fitness consequences of the behavior. An example could be tattooing in Polynesian cultures, highly valued in cultural context but viewed as excessive outside of it. Such excessive behaviors can lead to stigmatization outside of the group and therefore can create very strong group boundaries which facilitates the participation of group members in group activities as envisioned by Iannacone. This could then help groups deal with specific kinds of public goods problems. It also may facilitate norm enforcement through cheap punishment by increasing the consequences of exclusion from the group, as is part of the strategic effect of tattooing in Central American street gangs (Brenneman, 2011). Thus assuming rational actors, intent/behavior will be constrained by the structure of the stigma. Where conspicuous display of altruism itself is the trait biased toward, this can lead to runaway cultural evolution of generosity similar to the way it is envisioned by Gintis et al (2001), though this would eventually be limited by other constraints.

### Ethnic Markers and Shared Norms (McElreath, Boyd, & Richerson, 2003)

It is worth mentioning the theory of ethnic marking and shared norms as spelled out by McElreath et al‥ While it does not specifically model *costly* markers and it does not support *altruism* but *coordination*, it bears some similarities to these other theories of rituals as signals of group membership and is worth disambiguating. In this social learning model, it is assumed that people are faced with regular coordination problems. An example would be which side of the street to drive on or any sort of activity that depends on uniformity of decision amongst options, but where the specific option conformed to is irrelevant for payoff. In such a situation, there is no individual level incentive to ‘cheat’, but instead the dilemma of conforming to the same choice without prior communication. It is further assumed that 1) individuals preferentially engage in activities with individuals who are similarly marked and 2) individuals preferentially socially learn behaviors and markings together as packages with a bias based on payoff. Under such conditions, a diversity of coordination behaviors can coexist in the population and they become ‘marked’ … associated with some signal behavior, such as performing a ritual or dressing a certain way. It is thus similarly a theory of how ritual may evolve and be a relatively honest signal of intent. However, it does not solve public goods type problems, but only coordination problems, where group members are not required to be altruistic, but merely to coordinate. It is possible that some other mechanism may take advantage of such markings to foster altruism, but there would have to be some other social force at play. Also, it does not model *costly* signals. Of course, since it does show that such a psychology would be favored by evolution, it stands to reason that such markings could be costly to some degree and still be favored. However, such costs would be essentially efficacy costs and would not have a strategic function. Less costly markings would be favored when available.

Table 1 summarizes these various theories of cost in relationship to signaling.

**Table 1.** Theories of cost in relationship to signaling.

| Theory | Short description | Problem Solved | Effect of cost in model | Can solve public goods problem? Limits? | Reference |
| --- | --- | --- | --- | --- | --- |
| **Efficacy cost of manipulative signal** | The cost is necessary for idiosyncratic reasons in order to do something useful and is therefore supported. Includes indexical signals, where signal not mechanically possible without ability to pay cost. | None. One individual is manipulating another with a signal. Can be costly, but cost itself is not useful | Cost does not solve problem, but is justified as necessary to gain a benefit. Without cost, signal system works better. | No | (Guilford & Dawkins, 1991) |
| **Strategic handicap cost** | Only high quality individuals are capable of paying an obvious cost. Quality is important for receivers to know as they try to pick partners | Honest signal of one's higher quality to a potential partner | Cost paid by signaler allows sorting by receivers of signalers by quality. Without cost, signal system devolves. | No, although a conspicuous contribution to a public good may itself be a cost supported by this model, so it may in this case solve a public goods problem. | (Grafen, 1990; Zahavi, 1975) |
| **Stigmatizing costs** | Stigmatizing group practices force group members to participate in group activities. | Solves club good problems, where individuals would otherwise be able to take advantage of club goods while participating in beneficial exchanges outside of the group and not contributing to the group. | None, except insofar as costly activities may be stigmatizing. Stigma is what allows solution to club goods problem. Costless stigma would work as well. | Some. Only solves those PG problems where group members are forced to actively participate in costly group beneficial activities. Does not solve PG problems where individuals may choose to free-ride within groups. | (Iannaccone, 1992) |
| **hard to fake rituals** | The perceived cost or benefit of a ritual practice changes with true belief in a groups' religion. Where the religion also involves parochialism, this supports positive assortment amongst altruists | Problems of altruistic groups filtering out free-riders | Costs are essential for the filter. Without the cost, religions invaded by free-riders and true belief disappears from the population | Yes. Depends on how costly it is to fake relative to group benefits and the specific nature of the cost. Different potential interpretations produce different results and limits. | (Irons, 2001) |
| **Costly bonding practices** | Socially learned or genetically - based collaborative ritual practices hijack pre - existing genetically based social instincts to create altruistic bonds. | Problems of altruistic groups filtering out free-riders | Costs of ritual practice is sometimes unnecessary, sometimes allows for coexistence of free-riders and altruists. | Yes. Evolutionary stability of solution is dependent on relative costs/benefits of PG dilemma, hijacked gene, and ritual participation | (Frost, 2016a, 2016b) |
| **Runaway Social Learning** | Where there is a bias to socially learn a behavior and this bias is itself socially learned, this may lead to runaway cultural evolution of behaviors, including costly ones | How to stabilize costly behaviors in a population | Cost is not required by the model, but the dynamic may support a cost. | Not intrinsically although 1) conspicuous altruism may be a specific cost supported by the dynamic and 2) where such runaway processes lead to stigmatization of group members from greater society, it may help solve some specific public goods problems as stigmatizing costs. | (Boyd & Richerson, 1985) |
| **Ethnic Markers and Shared Coordination Norms** | As individuals are faced with coordination dilemmas, if individuals have a tendency to interact with those with similar markings and to pick up coordination behavior and markings as a package based on success biased social learning, then multiple | How to develop shared coordination norms within ethnically marked sub - populations | Costs are not explicitly modeled but could be tolerated to a limited degree as efficacy costs, though cheap markings would be favored. | No, only coordination problems | (McElreath et al., 2003) |

|  |  |  |  |  |  |
| --- | --- | --- | --- | --- | --- |
|  | coordination behaviors can coexist and they will be marked |  |  |  |  |
| **Credibility Enhancing Displays (CREDS)** | If we believe that a costly behavior is credibility enhancing for claims of having a belief and we believe that such CREDS are more credibility enhancing if they are more costly , we are more likely to socially acquire a true belief from someone claiming a belief if they also engage in costly CREDS, and costly CREDS will be supported in the population | How to maintain costly displays in a population. | Cost of display bolsters both associated belief and the display | No, the CRED psychology is proposed but not shown to evolve or be able to deal with free-riders. | (Henrich, 2009) |

## Part2: Modeling the hard-to-fake ritual marker theory

Before we can model the hard-to-fake model, we need to spell out more specifically what ‘cost’ and ‘fake’ mean. By ‘costly’, we take it that we have some sort of aversion to faking the signal or difficulty learning to do so. This could be taken to mean that it is something that we would never do, that is for some reason just not possible. It could mean that it is something that we are capable of doing, but which we find unpleasant or unattractive to do and so would only do if there was a strong incentive; we always know of it as an option and decide whether to do it or not based on an intuitive optimization of pleasure or ‘utility’. The aversion could instead manifest as a bias against socially learning to do the behavior. In this case, as we observe someone performing a ritual, we would be less likely to choose to copy them unless we also acquired the religious belief. To ‘fake’ the ritual would be to do it without having the complete set of religious or cultural group beliefs associated with the ritual. I’ll explore these interpretations separately in the following models.

All of the following are cultural evolution models, individuals social learning beliefs by copying peers. In longer time frames where there is genetic variance in social learning biases and there are costs and benefits measured in genetic fitness consequences, this may be relevant and we may need to use models of gene-culture coevolution (Frost, 2016b), but here I ignore such changes and just look at cultural evolution processes. Similarly, I focus on horizontal transmission of socially learned behaviors and do not look at dynamics of vertical transmission, which would entail tracking biological reproductive dynamics. It has been shown in the US that much of the change in the religious beliefs in the US in the last century is driven by different rates of biological reproduction, so this will certainly be important in some cases (Hout, Greeley, & Wilde, 2001), but I leave this for a future modeling exercise. The behavioral variants in the sub-models below are A1 (non-religious) and A2 (true believer). True believers perform visible rituals. In some of the models A1 may also fake rituals strategically. In other versions, faking the ritual is not a strategic decision, but one that is learned. In these models, there is a third behavioral variant, A3, that performs the ritual but does not have the prosocial behaviors of A2.

Individuals engage in Public Goods Games (PGG) with groups of others, where individuals contribute benefit b to the group by paying a cost c. The benefit is distributed evenly to all group members, so each group member receives b*r benefit, where r is the fraction of the group that chose to cooperate and pay the cost c. The benefit and cost are in terms of some experienced and observable utility, which in turn biases one’s efficacy as a mode for social learning. It leads to an increase or decrease in the probability of being copied by another. The most simple interpretation is that they are observable measures of some kind of culturally relevant form of success (like health, wealth, displayed happiness) and that social learning is ‘success biased’. In economics terms, we might call this success in acquiring additive contributions to utility.

Part of ‘true belief’ is a belief in cooperating with and altruistically sacrificing for other believers. Belief itself is not visible and thus can not directly be the basis of assortment. As we consider the possibility of performing rituals without true-belief, we need to consider our assumptions about social learning dynamics, since ritualists, by definition are advertising themselves as true believers. How can people socially learn to fake belief via ritual participation if no ritual practitioner is publicly displaying the fact that they do not believe? There are two extreme assumptions that could be made, and I explore both in the models below. The first is that anyone performing a ritual acts as an A2 (true believer) for social learning purposes. Anyone who sees them would learn from them to be a true believer, not a faker. The second possible assumption is that although fakers are effective at veiling their own lack of true belief, the existence of and fraction of non-believers amongst ritualists as well as the relative fitness benefits can be deduced. This deduction may happen from observed success amongst the group of ritualists as a whole, understanding of the implied public goods situation, and knowledge of the potential success of the group if all group members cooperated. With this second assumption, it is this set of observations and inferences that is used for social learning, not the expressed belief of group members. In this case, social learning is not from direct observation, but from inferred frequencies of behavior.

Talk is cheap and plays no explicit part in the public goods game and the models. While there may be some effects of cheap talk in the real world, it is not a part of these models. These models are of a more conservative assumption that such talk will have no effect.

Currencies of costs and benefits may be in utility maximization for some kind of optimization decision or may be in biasing of social learning, depending on the model. In the models below, when individuals are said to be able to do something “strategically”, this is defined as having multiple potential behaviors that the individual chooses from based on utility maximization. In other models, behavior is not strategic, but fixed once they are acquired through copying (social learning) or personal innovation. Where there is a potential for confusion, I clarify. While social learning is ‘success biased’ and *may* result in utility maximization at equilibrium, it is not the same thing as an individual engaging in utility maximizing decision making and at times certainly will not maximize utility.

p_A1_ is the population frequency of A1, p_A2_ is the population frequency of A2, and where there is an A3 variant, its frequency is given by p_A3_.

Success biased social learning assumes that when an individual meets another, they will copy the behavior of the other with a probability given by

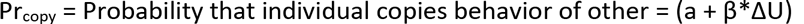

where a is the probability of unbiased copying, β is a measure of the strength of success bias, and ΔU is the difference between the observed markers of utility/success of the model and the experienced utility/success of the self. For the case where an individual chooses to either keep their behavior or adopt the behavior of the other completely randomly, with no success bias, we would have a = 0.5 and β = 0. a and β are constrained such that 0 ≤ Pr_copy_ ≤ 1. Where p_x_ is the frequency of X, the number of variant Y converting to variant Z in a given time step of social learning would be given by

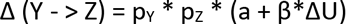

Equilibrium is found where the gross losses of cultural variants through social learning equal the gross gains. Pure populations are of course equilibriums. It is assumed that there is always introduction of variants through rare personal innovation. Pure populations are tested to see if they can be invaded, in order to find stable equilibriums.

Table 2 summarizes all parameter definitions. Table 3 summarizes the models analyzed below, with equilibrium results.

**Table 2.**
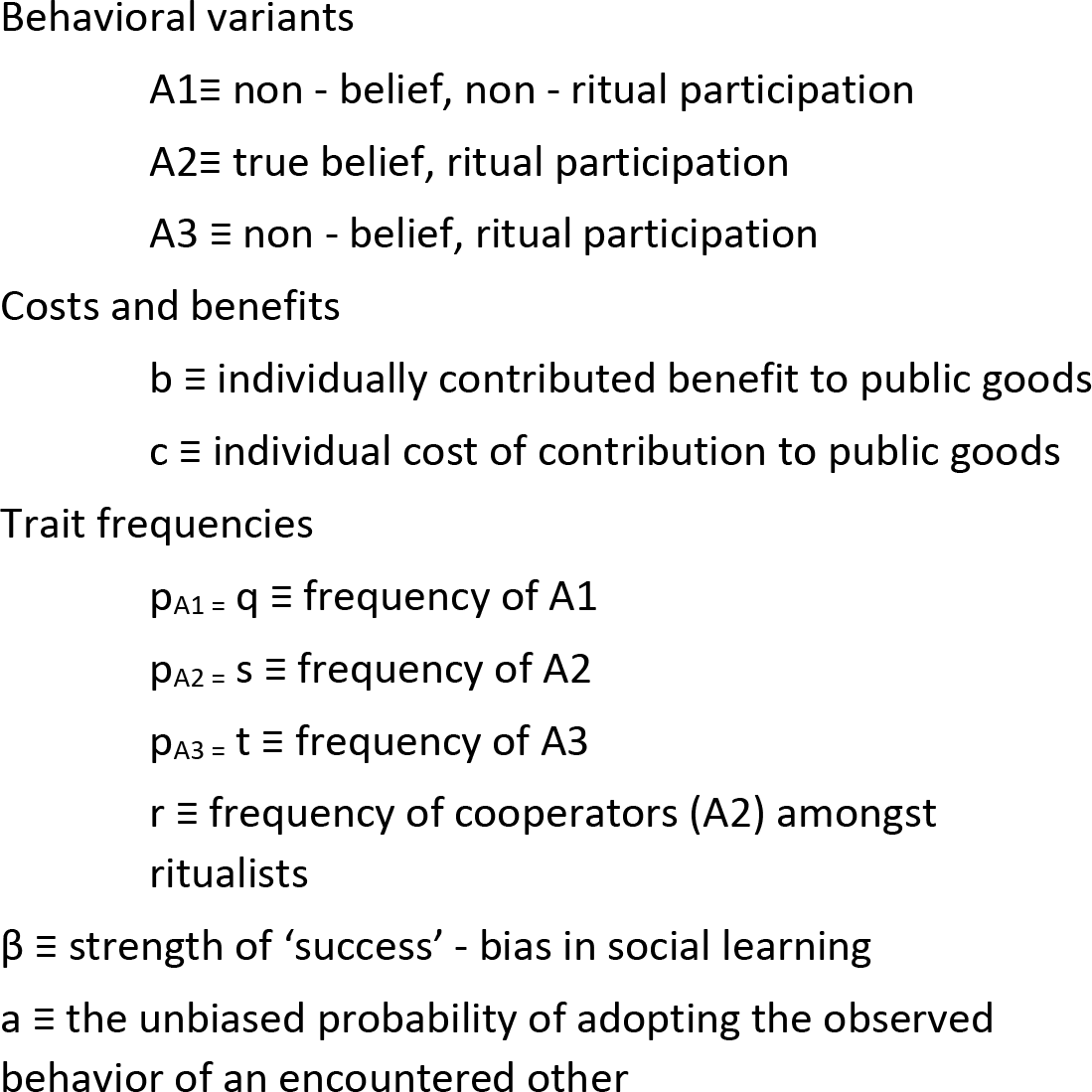
Parameters used in all models

## Model 1: “impossible to fake ritual”

‘Impossible-to-fake’ is the simplest interpretation of the hard-to-fake argument. Only true believers are willing to do the ritual. To non-believers, the ritual practice is too aversive to practice faithfully. Perhaps it is too painful, or perhaps the time investment is too heavy. Part of the belief system is a real commitment to altruism. It is not possible (due, for example, to cognitive limitations) to parse out the part of the beliefs that change one’s perspective on the ritual from the belief in altruism. The ritual, being an honest signal of true belief, can be relied on for weeding out free-riders. True believers can apply a simple heuristic of using the ritual as a marker of true belief, associating and cooperating only with other ritual performers as coreligionist. Ritual performers will be more strongly marked as ‘successful’ because of net benefit from mutual cooperation in the PD game. All other things being equal, they will dominate the population through success-biased social learning: 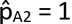.

Of course, this is the classic green beard solution to cooperation: if you have an honest signal of cooperation (‘all altruists have green beards’), that signal can be used to sort out free riders from cooperative altruists (Dawkins, 1976; Hamilton, 1964). A green beard solution works evolutionarily when there is no possibility of a green bearded free rider in the population. A variant which lacked such cognitive restraints and was able to perform the ritual without adopting the prosocial beliefs would be such a green bearded free-rider.

## Model 2: “Can strategically fake ritual, but we are averse to do so”

A1 in this model can strategically pretend to be religious and perform the rituals, but they are averse to doing so. We can quantify this aversion, f, in the same currency as b and c of the PGG.Non-believers experience this ritual cost. Believers do not. If an A1 becomes an A2, they no longer experience the cost, f, but are then committed to cooperative altruism.

Trivially, if f > b, then A1 never fakes the ritual. In this case, the cost is never worth the benefit. As A2 then always has a higher utility and social learning of beliefs is success biased, A2 again comes to dominate the population. This replicates the results of Model 1.

If f < b, then it is possible for some A1 individuals to join groups of A2s as free-riding, ritual-performing, signal-fakers. Ritual performers will always only associate with other ritual performers for the PGG. Since A1 individuals are utility maximizing, they will join groups of ritualists until the marginal benefit of doing so is zero (f = b*r) or until all A1 have become ritualists. This can be modeled as one ritual group: all A2s are part of this group and A1s join until it is no longer optimal to do so. The group members participate in a public goods game whose benefits are excludable to non-members. I assume that A1s rapidly join the group, but incrementally, observing the flow of benefits in the group, b*r. A1 individuals join the ritual group either until r = f/b or until all A1s are in the group, whichever comes first. r will be the greater of f/b or p_A2_. If p_A2_ < f/b, then there will be a sub-population of A1 individuals who do not join the group and do not perform rituals. U(A2) = (br - c). For p_A2_ ≤ f/b, only some A1 perform ritual, and U(A1) = 0. For p_A2>_f/b, all A1 perform ritual and U(A1) = (br − f).

We have established that the benefits and costs of the PGG are some visible contributions to utility, but what of the visibility of f, an experience of aversion? For simplicity, we will say that this aversion manifests visibly as a decrease in visible utility/happiness and has an effect on efficacy as a social model in proportion to its effect on utility maximization, in the same way as b and c.

We must pause for a moment to consider social learning. Non-believers are performing rituals. They are in a way modeling religious belief without being religious. We might then state that those who fake rituals (A1s in ritual groups) act for social learning purposes as true believers (A2), even though they do not cooperate in PGG. Let’s return to this (in Model 2B) after considering the more simply modeled alternative. In Model 2A, instead of looking at ritual behavior to determine the frequency of religious belief, individuals deduce the frequency of actual religious belief based on an understanding of potential net benefits of the PGG and observing actual flow of benefits in the group.

### Model 2A (A1s that fake the ritual act as A1 for social learning purposes)

Individuals effectively see the actual frequencies of A1 and A2 and socially learn these belief systems through success biased social learning, weighted by their actual frequencies in the population. While A1s successfully hide themselves as individuals within the ritual practicing group and can thus free-ride, their presence and collective frequency is recognized. This would be plausible, for example, when social learning happens over a longer period of time and people are able to observe failures of cooperative work without being able to name who failed to cooperate. Anyone who has spent time in a cooperative will be familiar with this dynamic. In this case, A1 would model non - religious belief not through direct advertisement, but through their anonymous impacts on the group’s well-being. The fitness of A1s, whether ritual performing or not, are all the same, otherwise, they would strategically shift to a higher fitness strategy.

The difference in visible success would be given by V (A2) − V (A1) = (f − c).

In short, for f > c, A2 takes over the population (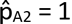) and for f < c, A1 takes over the population (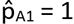). There is not a mixed equilibrium, except when f = c. The interpretation of this is pretty straightforward. If free-riding and its benefits are obvious in the population, it will come to dominate the population through success - biased social learning unless the intrinsic aversion to performing or learning the ritual practice for non - believers is greater than the potential advantage from free-riding. f > c gives the same result as Model 1. If f < c, then fakers will take over the group.

### Model 2B (A1s that fake the ritual act as A2 for social learning purposes)

In Model 2B, an A1 that performs ritual is treated as an A2 for social learning purposes. They successfully advertise themselves as true believers. When a social learner encounters a ritual-faking A1, what they learn from this A1 is true belief, A2. Thus, an A2 encountering such an A1 would not change its behavior, and an A1 encountering such an A1 might socially learn true belief, A2. However, what is visible is *its* utility, not the utility of an A2. It acts as a model of A2, but with the success biasing of a ritual-faking A1. The observed fitness of an A1 performing the ritual is its actual fitness, not the fitness of an A2, though what it models, deceptively, is A2.

A1 would never be able to invade a population of A2. In a world where religious belief was acquired purely through copying, no one would be advertising A1 to be copied. An invading A1, introduced perhaps through personal innovation, would be successful themselves themselves, but they would never serve as a model for social learning of A1, since they would be pretending to be A2.

In a population of A1s, on the other hand, when A2s are introduced, they would have an immediate boost in terms of social advertisement by A1s that joined their groups to freeride. Such A1s joining the group of A2s would appear to be A2s for social learning purposes. For f > c, this would just accelerate the inevitable domination of A2, so the equilibrium frequency is still 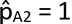. The frequency of A2 would appear higher than it actually is because freeriding A1s advertise themselves as A2s and act as A2 for social learning purposes. For f < c, even though A2s would have lower average ‘utility’, this would potentially be compensate for by the increased (false) advertising of A2 by free-riding A1s. In this case, there would be the illusion of a frequency of A2 given by p’_A2_ = (b/f) p_A2_. Whether the population was dominated by A1 or A2 would depend on the strength of success bias for social learning, with a strong success bias being necessary to overcome the illusion of more models of religious belief.

With this illusion in place, we have

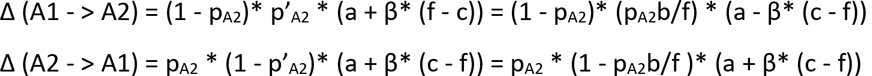

Now, if Δ (X) is the net increase in X, we have

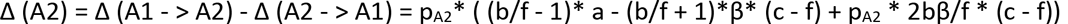

To see the conditions under which A2 can invade A1, we find as p_A2_ - > 0, Δ (A2) is positive for

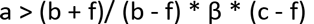

Moreover, if this condition is met, Δ (A2) is always positive. As anticipated, despite being invaded by cheating A1s such that the average utility of A2 is less than that of A1, A2 will successfully invade and take over the population because faking A1s represent themselves as A2s and therefore help in the social learning of A2. The conditions for this is set by the relative strengths of unbiased social learning vs success bias in social learning in relation to the other parameters. Alternately, when this condition is not met, the equation for Δ (A2) can be used to solve for p_A2,0_, the starting frequency of A2 that is necessary for A2 to be able to take off and take over the population

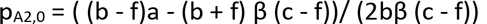

In short A1 can never invade A2, A2 can invade A1 and take over the population for a > (b + f)/ (b − f) * β * (c − f), and if, through some means, the population gets to p_A2_ > p_A2,0_, then A2 will take over the population, otherwise, if p_A2_ < p_A2,0_, then A1 will take over the population. There is a mixed equilibrium at p_A2_ = p_A2,0_, but it is unstable.

## Model 3: A3 is visible statistically but not individually. People averse to it

In model 4, people can see how well on average ritualists do. They ‘know’ or intuit b and c and therefore can somewhat tell r, the frequency of A2 amongst ritualists from how ‘well’ ritualists are doing. However, people have an intrinsic bias against learning A3. It could be that although they know somehow that there are free-riders, the religious belief system is hard to parse because of cognitive load or emotional constraints. In this model, there is a social learning bias against A3, given by f, again in the same currency as b and c, relative to success biased social learning.

Solving for equilibrium in social learning, we find for f < c, 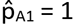. For f > c, 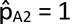. These are the same conditions and equilibrium frequencies as model 2A, where faking ritual is strategic, rather than socially learned through success biased social learning.

## Model 4: Individuals socially learn A3. A2 can strategically invest in distinguishing A2 from A3 individuals

A2 individuals would of course prefer to assort with other A2 individuals. In this model, they are strategically able to pay a cost in order to be able to figure out who is A2 and who is A3, which will become worth paying as r gets too low. Let’s call this the ‘free-rider filter’. Anyone who has ever lived in a cooperative house is familiar with this process and the associated costs. The dishes aren’t done and then you have to track down whose they are, everyone denies it, you become hypervigilant for leavers of dirty dishes (which is stressful) until you figure out who is doing it, and processing all of this is costly in terms of group conversations. This costly hypervigilance can payoff, though, in terms of eventually weeding out the freerider by kicking them out, converting them to an altruist, or getting them to behave altruistically through enforcement, now made cheap since you know who to look for. Then you have a period of trust and a release of the costly hypervigilance until turnover of roommates leads to too many freeriders again. A3 is not ‘costly’, but ritual is ‘hard-to-fake’ in that too many fakers causes the free-rider filter to be invoked.

The free-rider filter costs h to use. It functions to allow perfect assortment. When does it pay to use it? Assumedly, when the population of A3 is negligible, it doesn’t pay. If V’ is the fitness of the population when A2 switches to using the free-rider filter, then where

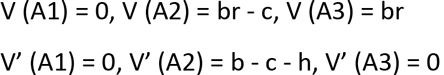

They switch to using the filter at V (A2) = V’ (A2), or r = (b - h)/b. For r > (b - h)/b, V(A2) > V’(A2), and it is better/cheaper for A2 to tolerate the free-riding A3s than to engage in costly filtering of free-riders.

Now, there are two possibilities. For h < b - c, A2 is effectively able to limit the relative frequency of free-riders amongst ritualists, and ritualists do consistently better than non - ritualists. Equilibrium frequencies are 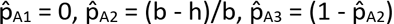.

There is something satisfying about this model in that we do feel that free-riders are an issue and yet groups persist and free-riders are limited. The possibility of investing in finding them will limit them, even if it is not constantly exercised. We might expect the population to oscillate between using and not using the freerider filter in response to rising and falling tides of A3.

In the case of h > b - c, as r < b/c, the A2 population socially learns to be non-religious (A1 or A3) through success-biased social learning and the population is left as a mixed population of A1 and A3, with A3 slowly increasing in frequency due to a vestigial population of A2’s, perhaps regularly re-introduced through innovation, to parasitize off of. 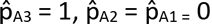

## Model 5: There is a flat cost to being in the religion (ritual performance), in addition to costly free-rider detection

As one final twist on costliness of ritual practice, assume there is a small, flat cost or aversion to being in the religion, given as a bias against socially learning to be in the religion, g. Unlike Model 4, this experienced cost is to everyone, believers and non - believers. We still have the possibility to invest in sorting out freeriders from true believers as in Model 5. We then have

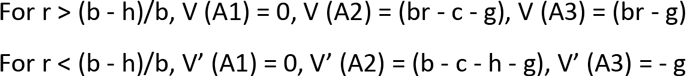

For h < (b - c - g), we have the same as h < (b − c), equilibrium frequencies are given by

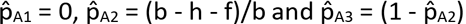

This parallels the results of model 5. However, for h > b - c - g, the cost of religious practice causes a qualitatively different end result than seen in Model 5. Here for r < (c + g)/b, A2s are disfavored by success biased social learning in comparison to both A1 and A3. The cost, g, causes staying in the religion to be less appealing for A3 as r < g/b, and A3 will also decline in tandem with A2. A3s will convert through success - biased social learning to A1 as altruistic A2s decline in frequency. At equilibrium, the population is entirely A1. This result appears to be consistent with the regularly recurring decline of communes as in-group cooperation declines(Sosis & Bressler, 2003). The flat cost does not save the ritual group, but can be seen as a way of preventing the persistence of groups of ritual fakers. In a population where A2 (true belief) is regularly re-introduced, this could allow for regularly recurring populations of true believers that eventually get eliminated by free riding A3s, who then eventually convert back to A1(Jansen & van Baalen, 2006).

## Conclusions

These are simple models and undoubtedly real world dynamics are more complex, and a more realistic model would contain a combination of these dynamics. Still, these simple models suggest the qualitatively dynamics one might observe, where conditions formalized in the models apply. The premise of the theory of ritual as costly-to-fake signal of true-belief and altruism as proposed by Irons and explored empirically by Sosis and other is found to be cogent under a wide variety of interpretations, but specifics of what it might mean for the ritual to be ‘costly’ or ‘hard’ to fake create limits on when the logic holds. In looking at the results of the model analyses, first, an obvious conclusion comes out of the models that where the aversion to faking the ritual is greater than the cost of cooperation in the Public Goods Game (f < c), such ritual fakers are successfully weeded out of the population and ritual-marked cooperation flourishes. This holds as a condition for ritual facilitated altruism to take over the population whether this cost of faking is an experienced aversion for a strategic decision to fake or an aversion to socially learn to fake the ritual. Second, the analysis uniquely looks at cases of successful false-advertising and finds that such can actually be advantageous to religious groups, despite free-riding. Third many of the models support the conclusions of the hard-to-fake ritual theory, but present conditions that must be met for such hard to fake rituals to effectively stop free-riders from taking over a population, and in some of these cases, equilibrium is characterized by a balance of free-riders and true believers performing ritual.

The case of people performing behaviors that they are not advertising sets up an interesting conundrum for the perspective of social learning models of behavioral change. For the more extreme case of completely successful false advertising, where those faking the ritual successfully represent themselves as true believers and thus successfully model and teach true belief in social learning, the disadvantages of free-riding can be compensated for with the help that such free-riders inadvertently offer in social learning. Such fakers still bring in converts. Specifically, the model (2B) predicts that under some parameter combinations true believers will always take over the population and that for others, they will still take over the population if they somehow get above a threshold frequency in the population. If some circumstance leads to this starting frequency, A2 will take off through social learning boosted by A1s pretending to be A2, A2 will dominate the population, and 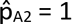. The trajectory of such a population is quite interesting. At first the religion is full of fakers and does not do well in terms of delivering satisfaction to members, but because there is an illusion of people joining and adhering to the beliefs more than they actually are, the religion takes off and the fakers themselves actually disappear from the population, as they have the impression that they are alone and eventually give up their non-belief (despite the utility advantage) because of social learning pressures. This occurs without hyper-conformity, punishment, or ostracism of non-cooperators. If this were an accurate modeling of the social learning dynamics, religious leaders could strategically modulate isolation of members, opening up if p_A2_ > p_A2,0_, to attract members and becoming more closed off if p_A2_ < p_A2,0_, in order to artificially create an environment where p_A2_ > p_A2,0_, and thereby bolster commitments to belief within the religion, or maintain the effective frequency of their members in the sub-population above p_A2,0_ to ensure expansion of the group. Similarly, other more costly methods of bolstering group membership might be utilized until p_A2_ > p_A2,0_, at which point, cultural evolution would carry true belief to fixation without costly investments. Of course non-believers could never invade a population of true believers in this model. Any non-believer introduced would model true belief and they would eventually convert without causing any apostasy. If non-believers invade, it must be through some other mechanism.

**Table 3.**
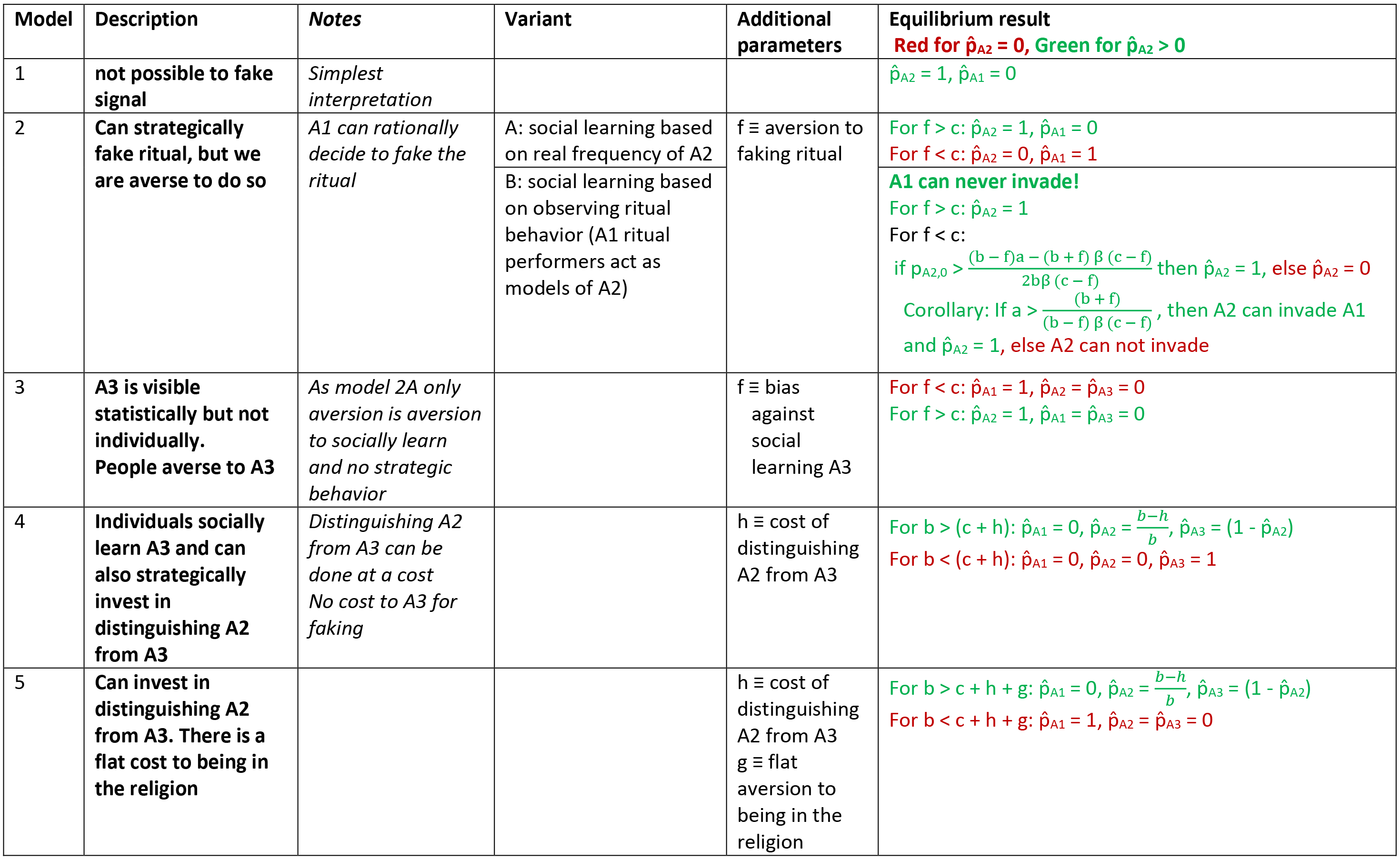
Models implementing different interpretations of the costly-to-fake signal theory

| Model | Description | Notes | Variant | Additional parameters | Equilibrium result<br>**Red for $\hat{p}_{A2} = 0$, Green for $\hat{p}_{A2} > 0$** |
| --- | --- | --- | --- | --- | --- |
| 1 | not possible to fake signal | *Simplest interpretation* | | | $\hat{p}_{A2} = 1, \hat{p}_{A1} = 0$ |
| 2 | Can strategically fake ritual, but we are averse to do so | *A1 can rationally decide to fake the ritual* | A: social learning based on real frequency of A2 | $f \equiv$ aversion to faking ritual | For $f > c$: $\hat{p}_{A2} = 1, \hat{p}_{A1} = 0$<br>For $f < c$: $\hat{p}_{A2} = 0, \hat{p}_{A1} = 1$ |
| | | | B: social learning based on observing ritual behavior (A1 ritual performers act as models of A2) | | **A1 can never invade!**<br>For $f > c$: $\hat{p}_{A2} = 1$<br>For $f < c$:<br>if $p_{A2,0} > \frac{(b-f)a - (b+f)\beta(c-f)}{2b\beta(c-f)}$ then $\hat{p}_{A2} = 1$, else $\hat{p}_{A2} = 0$<br>Corollary: If $a > \frac{(b+f)}{(b-f)\beta(c-f)}$, then A2 can invade A1 and $\hat{p}_{A2} = 1$, else A2 can not invade |
| 3 | A3 is visible statistically but not individually. People averse to A3 | *As model 2A only aversion is aversion to socially learn and no strategic behavior* | | $f \equiv$ bias against social learning A3 | For $f < c$: $\hat{p}_{A1} = 1, \hat{p}_{A2} = \hat{p}_{A3} = 0$<br>For $f > c$: $\hat{p}_{A2} = 1, \hat{p}_{A1} = \hat{p}_{A3} = 0$ |
| 4 | Individuals socially learn A3 and can also strategically invest in distinguishing A2 from A3 | *Distinguishing A2 from A3 can be done at a cost*<br>*No cost to A3 for faking* | | $h \equiv$ cost of distinguishing A2 from A3 | For $b > (c+h)$: $\hat{p}_{A1} = 0, \hat{p}_{A2} = \frac{b-h}{b}, \hat{p}_{A3} = (1 - \hat{p}_{A2})$<br>For $b < (c+h)$: $\hat{p}_{A1} = 0, \hat{p}_{A2} = 0, \hat{p}_{A3} = 1$ |
| 5 | Can invest in distinguishing A2 from A3. There is a flat cost to being in the religion | | | $h \equiv$ cost of distinguishing A2 from A3<br>$g \equiv$ flat aversion to being in the religion | For $b > c+h+g$: $\hat{p}_{A1} = 0, \hat{p}_{A2} = \frac{b-h}{b}, \hat{p}_{A3} = (1 - \hat{p}_{A2})$<br>For $b < c+h+g$: $\hat{p}_{A1} = 1, \hat{p}_{A2} = \hat{p}_{A3} = 0$ |

In most models analyzed here, either true belief takes over the population or is completely eliminated from the population. The exception is where there can be a strategic investment in searching out and identifying those who are not true believers amongst ritual performers. This then creates a dynamic where either everyone performs ritual and there is balance of true believers and fakers or where no one performs ritual. In the case where filtering out free-riders is sufficiently cheap (h < (b-c)), the ratio of free-riding fakers to true believers can be kept sufficiently low and ritual performance takes over the population. The population would oscillate between filtering and not filtering as the population of fakers reciprocally rose and fell. If the cost of filtering is too high, true belief is eliminated from the populations (model 4), but it takes a small flat cost added to being in the religion, some small sacrifice, in order to actually eliminate ritual performance (model 5). One would imagine the same if the small cost was something that only fakers had to pay (as for example combining Models 4 and 2A).

Many of these models suggest phenomena that feel familiar for religions and may provide an underlying explanation. For example strategic isolation may be functional in terms of either negotiating proselytizing success or management of free-riders through reduction in models of non-belief (as suggested by the dynamics in Model 2B). The persistence, but non-domination of free-riders might be explicable in terms of the potential use of more hypervigilant searches for free-riders which may not currently be in use (Model 4). Persistence of altruists in groups dominated by free-riders might be explicable in terms of social learning and the false advertising of altruism by said free-riders (Model 2B); altruism may persist in such groups due to deceptive free-riders teaching altruism.

Next steps would be to examine real world religions or ritual-using groups for patterns of altruism in relationship to ritual costliness that might fit the predictions of these models. Specifically, it would be interesting to ask in future research…

- How pervasive are the dynamics of contextual investment in rooting out regularly recurring free-riders?
- What is the relative efficacy of non-believing free-riders in modeling altruistic true belief? Do deceptive non-altruists effectively teach altruism?
- What are people’s assumptions about the presence of hidden free-riders in groups, how and when do they invest in determining this, and, alternatively, do they socially learn free-riding through such deductions?
- What are the factors that make it easier or harder to break apart belief systems and learn only parts vs the whole package?
- What are the dynamics of cost that either make a failing communal group dissolve or become a stable grouping of non-cooperating free-riders?
- How does true belief shift the experience of otherwise dysphoric ritual practices? Is the dysphoria turned into ecstasy, or is the dysphoria balanced by a sense of obligation and fear? Is there a difference between ritual motivated by positive vs negative emotions in terms of ease or difficulty in faking?
- When someone converts to a religion in order to take advantage of the public good benefits of the group (as Sosis argued happened with changing economic circumstances with the Shakers over their 200 year history (Sosis, 2000)), what are the factors that control whether they actually adopt true belief or simply ‘fake’ the rituals, tolerating the dysphoria and leaving the religion as economic conditions outside of the religion improve?
- Similarly, as a state provides public goods, it potentially undermines religious groups’ attractiveness in providing public goods to its members. A number of these models suggest that this would cause religious fakers to leave religious groups because of the costly ritual practices. Numbers would decline, but quality of membership might go up. This presents a testable hypothesis of these models.
- There are other theories of how some kinds of costly activities might promote group cohesion. Examples include Whitehouse’s dual modes hypothesis of the direct emotional response of bonding from certain kinds of dysphoric rituals or McNeil’s suggestion of the instinctual bonding effects of synchronized movement. As we observe seemingly costly ritual activities and sacrifice in groups, there are first empirical questions of whether they help bond groups into functional cooperative units. Secondly however, there remains the question of which specific mechanism is at play. It will be an important empirical question to ask whether a specific ritual is a trigger of instinctual prosocial response, or whether its experience is transformed by true-belief and it therefore becomes a reliable marker of true belief and thus altruistic intent. It will be important to start to distinguish what rituals are using what mechanisms to facilitate prosociality.

